# Evolution and coexistence in response to a key innovation in a long-term evolution experiment with *Escherichia coli*

**DOI:** 10.1101/020958

**Authors:** Caroline B. Turner, Zachary D. Blount, Daniel H. Mitchell, Richard E. Lenski

## Abstract

Evolution of a novel function can greatly alter the effects of an organism on its environment. These environmental changes can, in turn, affect the further evolution of that organism and any coexisting organisms. We examine these effects and feedbacks following evolution of a novel function in the long-term evolution experiment (LTEE) with *Escherichia coli*. A characteristic feature of *E. coli* is its inability to consume citrate aerobically. However, that ability evolved in one of the LTEE populations. In this population, citrate-utilizing bacteria (Cit^+^) coexisted stably with another clade of bacteria that lacked the capacity to utilize citrate (Cit^−^). This coexistence was shaped by the evolution of a cross-feeding relationship in which Cit^+^ cells released the dicarboxylic acids succinate, fumarate, and malate into the medium, and Cit^−^ cells evolved improved growth on these carbon sources, as did the Cit^+^ cells. Thus, the evolution of citrate consumption led to a flask-based ecosystem that went from a single limiting resource, glucose, to one with five resources either shared or partitioned between two coexisting clades. Our findings show how evolutionary novelties can change environmental conditions, thereby facilitating diversity and altering both the structure of an ecosystem and the evolutionary trajectories of coexisting organisms.

*Evolution does not produce novelties from scratch. It works on what already exists, either transforming a system to give it new functions or combining several systems to produce a more elaborate one.*

*–Francois Jacob*

## Introduction

Although ecology and evolutionary biology are often treated as distinct disciplines, scientists as far back as Darwin have recognized that the two are closely intertwined (Darwin 1881; Dawkins 1982; Lewontin 2001). Recent research on the interplay between ecological and evolutionary processes (Matthews et al. 2011; Schoener 2011) has emphasized that the two can interact even over short time scales. Evolutionary innovations—and in particular the emergence of qualitatively new functions—have the potential to alter ecological conditions both for the organisms that evolved the innovation and for coexisting organisms. Perhaps the most dramatic example of this interplay in the history of life on Earth was the evolution of oxygenic photosynthesis. This new way of obtaining energy from light was extraordinarily successful, but it produced oxygen as a by-product. Over time, oxygen accumulated in the atmosphere, where it acted as a toxin to most extant life (Knoll 2003; Sessions et al. 2009). Many organisms were likely driven extinct, but others evolved mechanisms to tolerate and use oxygen; thus, the oxygenated world created new ecological niches requiring aerobic respiration and perhaps favoring multicellularity (Schirrmeister et al. 2013).

The origin of a novel function is, by its nature, a rare event. Examples such as the evolution of oxygenic photosynthesis can only be studied using the fossil and geological records, making it difficult to resolve the order of events and examine the consequences of the innovation. The evolution of a new function can lead to major changes in an ecosystem, as some lineages may be driven extinct while others evolve into new niches. Ideally, one would like to be able to study the ecological and evolutionary consequences of an evolutionary innovation by directly comparing the properties of living organisms from before and after the innovation and measuring their relative fitness by competing the ancestral and derived forms in the environments where they evolved. In fact, the evolution of a novel function in a laboratory evolution experiment gives us the opportunity to do just that.

The long-term evolution experiment with *Escherichia coli* (LTEE), in which this novel function arose, is an ongoing experiment in which 12 populations founded from a common ancestor have been evolving in identical environments for more than 25 years. The initial ecology of the LTEE was deliberately made very simple, with the bacteria growing on a single carbon source, glucose, in a carbon-limited medium (Lenski et al. 1991). The culture medium also contained a large quantity of a potential second carbon source, citrate, but this resource was not available for *E. coli* metabolism. In fact, the inability to grow aerobically on citrate is one of the diagnostic features of *E. coli* (Koser et al. 1924). However, after more than 30,000 generations, one of the populations, called Ara-3, evolved the ability to use the citrate (Blount et al. 2008). After 33,000 generations, the citrate-consuming bacteria (Cit^+^) became numerically dominant in the population, and the total population size increased several fold. Following this large increase in numbers, much of the existing diversity in the population was eliminated, but a clade that could not consume citrate (Cit^−^) persisted and exhibited negative frequency-dependent coexistence with the Cit^+^ cells (Blount et al. 2008; Blount et al. 2012). The clade of Cit^−^ cells that persisted and the clade in which the Cit^+^ phenotype arose had diverged about 10,000 generations before the origin of the Cit^+^ function. All of the Cit^−^ cells isolated after the Cit^+^ type became numerically dominant came from this same clade (Blount et al. 2012).

We set out to study the ecological consequences of the evolution of citrate consumption and how it affected the further evolution of both the Cit^+^ and Cit^−^ lineages. In particular, we focused on studying the mechanism by which the Cit^−^ clade coexisted with the Cit^+^ clade. At least two distinct mechanisms could underlie this coexistence. First, a tradeoff between growth on glucose and citrate might allow the Cit^−^ lineage to persist as a specialist on glucose. Indeed, a comparison of the growth curves of Cit^+^ and Cit^−^ cells showed that the Cit^+^ cells had a longer lag time when growing under the same conditions as used in the LTEE (Blount et al. 2008). Second, Cit^−^ cells might cross-feed on one or more carbon compounds released by the Cit^+^ cells. We define cross-feeding as the consumption by one group of organisms of products produced by another group of organisms. The two mechanisms are not mutually exclusive and could act together to enable the coexistence of the Cit^+^ and Cit^−^ lineages.

The emergence of cross-feeding interactions has been observed repeatedly in evolution experiments with *E. coli* (Rosenzweig et al. 1994; Treves et al. 1998), and it is predicted to occur by theory under certain conditions (Doebeli 2002). Cross-feeding occurs when, in the process of growing on a primary resource, one bacterial ecotype secretes a byproduct that is consumed by another ecotype. Cross-feeding is favored when bacteria can grow more quickly by not fully degrading their primary resource, but instead secrete incompletely digested byproducts, opening the opportunity for the evolution of specialists on the byproducts. Cross-feeding has evolved in at least one other population of the LTEE, in which two lineages have stably coexisted for tens of thousands of generations (Rozen and Lenski 2000; Le Gac et al. 2012); in another population, a negative frequency-dependent interaction arose, probably also involving cross-feeding, that led to transient coexistence for several thousand generations (Maddamsetti et al. 2015; Ribeck and Lenski 2015). However, cross-feeding interactions may be less common in the LTEE than in some other evolution experiments because the relatively low glucose input and resulting low bacterial density limit the accumulation of secondary metabolites (Rozen and Lenski 2000). Moreover, mathematical modeling shows that cross-feeding will be less favored under a serial-transfer regime like that of the LTEE, where population densities are low much of the time, than in continuous culture, where populations are always near their maximum density (Doebeli 2002).

One potentially important set of molecules for cross-feeding in the Cit^+^/Cit^−^ system includes the C_4_-dicarboxylic acids succinate, fumarate, and malate. The Cit^+^ phenotype is based on the aerobic expression of the gene encoding the CitT transporter protein (Blount et al. 2012). CitT is a generalized di- and tricarboxylic acid antiporter that can import citrate, succinate, fumarate, and malate in exchange for export of any of these molecules (Pos et al. 1998). Thus, any net import of citrate into the cell requires that the CitT transporter pump succinate, fumarate, or malate out of the cell, making these C_4_-dicarboxylates likely molecules for cross-feeding.

Here we show that a key innovation in a long-term experiment with *E. coli*, the evolution of aerobic citrate metabolism, altered the ecological conditions experienced by the bacteria, influencing the subsequent evolution of both the lineage that evolved the new function as well as another surviving lineage that did not. In particular, we demonstrate that Cit^+^ cells modified the environment by releasing additional carbon sources—the C_4_-dicarboxylates—into the medium. The coexisting Cit^−^ bacteria evolved to cross-feed on those resources, adapting to the modified environment but resulting in a cost to their fitness when growing on glucose in the absence of the Cit^+^ cells. The Cit^+^ bacteria also evolved improved growth on the additional resources, thus competing with the Cit^−^ lineage for both the glucose and C_4_-dicarboxylates.

## Methods

### Long-term evolution experiment with *E. coli*

The long-term evolution experiment (LTEE) consists of 12 evolving populations of *E. coli* B, which are serially propagated in 10 mL of Davis minimal medium with 25 mg/L glucose (DM25). All populations were initiated from two founder clones that differ by a neutral mutation. Cultures are grown in a 37°C shaking incubator under well-mixed, aerobic conditions. Growth in DM25 is carbon-limited with a final population density of ∼5 × 10^7^ cells/mL in the absence of citrate consumption. Growth of Cit^+^ cells remains carbon-limited (Fig. S1) with a population density of ∼2 × 10^8^ cells/mL. Each population is diluted 1:100 into fresh medium daily, allowing regrowth of the population for ∼6.6 generations per day. Every 500 generations, samples of each population are frozen at -80°C. These frozen organisms are viable and can be revived for later experimentation. More details on the methods used in the LTEE are given in Lenski et al. (1991).

### Strains used and conditioning procedure

Other than the ancestral clone, REL606, all clones were isolated from the LTEE population, designated Ara-3, in which the ability to consume citrate evolved. Unless otherwise specified, the Cit^−^ clones used in this study were from the Cit^−^ clade (called clade 2 in Blount et al. 2012) that persisted after the evolution of citrate consumption. Clones from the Cit^+^ clade vary in their ability to grow on citrate. All clones from the Cit^+^ clade (called clade 3 in Blount et al. 2012) used in this study that were isolated at 32,000 generations or later have a Cit^+^ phenotype; however, clones from that clade that were isolated at earlier time points have a Cit^−^ phenotype. Members of the Cit^−^ and Cit^+^ clades were identified by mutations unique to each lineage, as described in Blount et al. (2012). Table S1 lists all of the strains used in this study.

Prior to initiating each experiment, we revived samples of frozen isolates by first growing them in Luria-Bertani broth (LB). We then conditioned them to the environment of the LTEE by growing them for 2 days in DM25, with transfers into fresh medium every 24 h.

### Resource assays

We hypothesized that the Cit^+^ cells released C_4_-dicarboylates into the culture medium, as a consequence of the antiporter mode of action of the CitT protein. To test this hypothesis, we employed gas chromatography–mass spectrometry (GCMS) to measure the concentrations of succinate, fumarate, and malate in the medium over the course of a 24-h transfer cycle. We measured these C_4_-dicarboxylate concentrations in cultures of the following strains: REL606, the founding strain of the Ara-3 population; a 40,000-generation clone from the Cit^−^ clade; and eight clones from the Cit^+^ clade sampled every 2,000 generations from 30,000 to 44,000 generations. Of the clones from the Cit^+^ clade, the 30,000-generation clone had a Cit^−^ phenotype, the 32,000-generation clone exhibited weak growth on citrate, and the remaining clones exhibited strong growth on citrate. After reviving and conditioning the clones as described above, we transferred the cultures to fresh DM25 medium. At 0, 2, 4, 6, 8, 12 and 24 h after transfer, 300 mL of each culture was collected using a sterile syringe, immediately filter-sterilized by passing the culture medium through a 0.25-μm filter, and frozen at -80°C for later analysis.

We measured the succinate, fumarate, and malate concentrations in these samples using an Agilent 5975 GCMS system at the Michigan State University Research Technology Support Facility. Because DM25 medium contains a high concentration of phosphate that obscured all other molecules in full-scan mode, we could not conduct a broad chemical survey. We instead used selective ion measurement to isolate the spectral signatures specific to succinate, fumarate, and malate. Because of this limitation, we cannot eliminate the possibility that Cit^+^ cells also released other molecules into the culture medium.

We also measured the concentration of citrate in the medium to compare the timing of citrate consumption to the timing of succinate, fumarate, and malate release. We measured citrate consumption by the 30,000-, 32,000-, 34,000- and 40,000-generation Cit^+^ clones with five-fold replication. We collected filtered samples of culture medium at 0, 1, 2, 3, 4, 6, 9, and 24 h, following the same protocol as above. We then measured the concentration of citrate in these samples using a Megazyme citric acid assay kit.

### Growth using C_4_-dicarboxylates as carbon sources

To assess changes in the ability of clones from the Cit^−^ and Cit^+^ clades to grow on succinate, fumarate, and malate, we analyzed growth curves for Cit^−^ and Cit^+^ clones isolated every 1,000 generations from 30,000 to 43,000 generations. This range includes clones from three periods of interest: (i) prior to the origin of the Cit^+^ phenotype (30,000 to 31,000 generations; (ii) when Cit^+^ cells were present, but very rare (32,000 to 33,000 generations); and (iii) after the several-fold population expansion of the Cit^+^ lineage (34,000 to 43,000 generations). The growth trajectories of each clone were assessed for three replicate cultures in media with a single alternative carbon source in the place of glucose. In addition to the C_4_-dicarboxylates succinate, fumarate, and malate, we also measured the growth of the Cit^−^ clones on acetate, a common substrate for cross-feeding in *E. coli* (Rosenzweig et al. 1994; Treves et al. 1998).

We used glucose as a carbon source during conditioning so that all cultures would have a similar initial density. For the Cit^−^ clones, growth curves were conducted in DM medium. However, growth curves of the Cit^+^ clones were conducted in M9 minimal salts medium (Sambrook and Russell 2001) because DM medium contains citrate, which would confound the assays. As a result, the growth curves for the Cit^+^ and Cit^−^ clones are not directly comparable; instead, comparisons of the growth curves provide information on the changes that evolved within each clade over time. The concentrations of the carbon sources used were as follows: 30.5 mg-succinate/L, 39.5 mg-fumarate/L, 45.7 mg-malate/L, and 34.4 mg-sodium acetate/L. We chose these concentrations because they give stationary-phase population densities similar to those in DM25. Growth curves were generated in 96-well microplates by measuring optical density (OD) at 420 nm every 10 min for 24 h (or 48 h, where noted) using a VersaMax automated plate reader (Molecular Devices).

We analyzed two characteristics of the growth curves: the final OD value at 24 h, and the time required to reach stationary phase. We determined time to stationary phase by visually inspecting each log-transformed growth curve and identifying the time point when a population switched from exponential growth to either a stable or declining OD. These visual assessments were performed twice, in a blind fashion, and the results were extremely consistent between repetitions.

### Fitness assays

To determine how evolutionary changes in the Cit^−^ lineage affected the fitness of Cit^−^ cells in the presence and absence of Cit^+^ cells, we conducted a series of competition experiments between a reference 30,000-generation Cit^−^ clone and Cit^−^ clones sampled every 1,000 generations from 30,000 to 40,000 generations. All Cit^−^ clones used in these assays were from the Cit^−^ clade. All of the Cit^−^ clones competed against the reference clone, in both the presence and absence of a mutant derived from a Cit^+^ clone from 40,000 generations, with three replicates of each competition. Competitions ran for one standard 24-h cycle, and fitness values were calculated as the ratio of realized population growth rates during the competition assay (Lenski et al. 1991). The Cit^−^ clones used were the same as those for the growth measurements above, with the exception of 33,000-generations, for which we inadvertently used a different clone, CZB195, instead of CZB194. However, we confirmed in separate measurements that CZB195, like CZB194 was unable to grow on the C_4_-dicarboxylates.

The reference clone, called CBT1, was an arabinose-consuming (Ara^+^) mutant of a Cit^−^ clone isolated from the 30,000-generation population (Table 1). Arabinose consumption is a neutral marker under the conditions of the LTEE (Lenski et al. 1991) that allows the differentiation of bacteria by their colony color when grown on tetrazolium arabinose (TA) agar plates (Levin et al. 1977; Lenski et al. 1991). To isolate CBT1, we grew the 30,000-generation Cit^−^ clone in 10 mL of DM25 medium, then centrifuged the culture and plated the concentrated cells onto a minimal arabinose agar plate. After incubation, a single colony from the plate was streaked twice on minimal arabinose agar plates. A colony from the second plate was grown in LB, and a sample of that culture was frozen. We then confirmed that CBT1 was selectively neutral compared to its parent clone by conducting a seven-day competition between the two clones under the same conditions as the LTEE.

To conduct competitions between Cit^−^ clones in the presence of Cit^+^ cells, we needed a way to prevent the Cit^+^ bacteria from growing on the TA plates. To this end, the competitions were conducted using a λ-sensitive mutant, called CBT3, of a Cit^+^ clone isolated from the population at 40,000 generations. Although the founding clone of the LTEE was sensitive to bacteriophage λ, resistant mutants were favored and went to fixation in population Ara-3 by 10,000 generations (Meyer et al. 2010). Resistant strains exhibit reduced expression of LamB, a maltose transport protein that is also the adsorption site for λ (Pelosi et al. 2006; Meyer et al. 2010). To isolate the λ-sensitive mutant, CBT3, we selected for a maltose-consuming mutant following the procedure described above for isolating an Ara^+^ mutant, substituting minimal maltose for minimal arabinose agar plates. We confirmed that CBT3 was sensitive to phage λ by plating onto TA plates with and without λ. The fitness of the Cit^+^ λ-sensitive mutant was similar to that of its λ-resistant parent in one-day competition assays (relative fitness 0.98 ± 0.06, mean ± 95% confidence interval, n = 10), and the Cit^+^ population size was unchanged.

For all competitions between Cit^−^ clones that were performed in the presence of Cit^+^ cells, the cultures were plated together with phage λ to prevent the growth of Cit^+^ colonies. However, a few Cit^+^ colonies were able to grow on these plates, presumably due to mutations to λ-resistance or incomplete coverage of λ on the plates. To correct for this problem, we transferred a sample of all Ara^−^ colonies onto Christensen’s citrate agar, an indicator medium for the Cit^+^ phenotype (Reddy et al. 2007). Any colonies that showed a positive reaction on Christensen’s agar were excluded from the counts used to measure the relative fitness of the Cit^−^ competitors.

Based on the outcome of these experiments, we decided to compete Cit^−^ clones isolated at 33,000 and 34,000 generations directly against one another. We measured the relative fitness of the 33,000- and 34,000-generation clones with ten-fold replication in both the presence and absence of Cit^+^, following the same procedure as above. Half of the replicates had the 33,000-generation clone competing against an Ara^+^ mutant of the 34,000-generation clone (named CBT2, isolated as described above), and half had the 34,000-generation clone competing against an Ara^+^ mutant of the 33,000-generation clone (called CZB207).

### Genomic analysis

The genomes of seven clones from the Cit^−^ clade were sequenced by Blount et al. (2012). We inspected the genome sequences to look for candidate mutations that might have affected the evolution of C_4_-dicarboxylate consumption. Three clones—ZDB357, ZDB200, and ZDB158 from generations 30,000 to 32,500 (Table S1)—could not grow on the C_4_-dicarboxylates. The other four clones—ZDB87, ZDB99, ZDB111, and REL10988 from generations 36,000 to 40,000 (Table S1)—exhibited strong growth on the C_4_-dicarboxylates. We used the breseq 0.23 software (Deatherage and Barrick 2014) to generate a list of mutations that were present in the C_4_-dicarboxylate-consuming clones, but absent in the clones that lacked the ability to grow on C_4_-dicarboxylates.

### Genetic manipulation

We constructed an isogenic derivative of the ancestral clone REL606 that carried the *dcuS* allele found in a 34,000-generation Cit^−^ clone called ZDB86. To do so, we used the pKOV plasmid and the methods in Link et al. (Link et al. 1997). We performed Sanger sequencing to confirm that the evolved *dcuS* allele was present in the resulting construct, ZDB1052.

## Results

### Resource assays

Consistent with our expectation based on the antiporter mechanism used for citrate uptake, we found that Cit^+^ clones secrete succinate, fumarate, and malate into the culture medium. The concentrations of these C_4_-dicarboxylates remained low throughout the 24-h transfer cycle in cultures of the ancestral strain (REL606), the 40,000-generation Cit^−^ clone, and a 30,000-generation clone from the clade that later gave rise to the Cit^+^ lineage (Fig. 1). By contrast, both succinate and fumarate concentrations were elevated in all cultures of all of the Cit^+^ clones during at least part of the 24-h cycle (Fig. 1A-B). Malate concentrations were also noticeably elevated in all except one (from generation 36,000) of the Cit^+^ clones that we analyzed (Fig. 1C).

**Figure 1.**
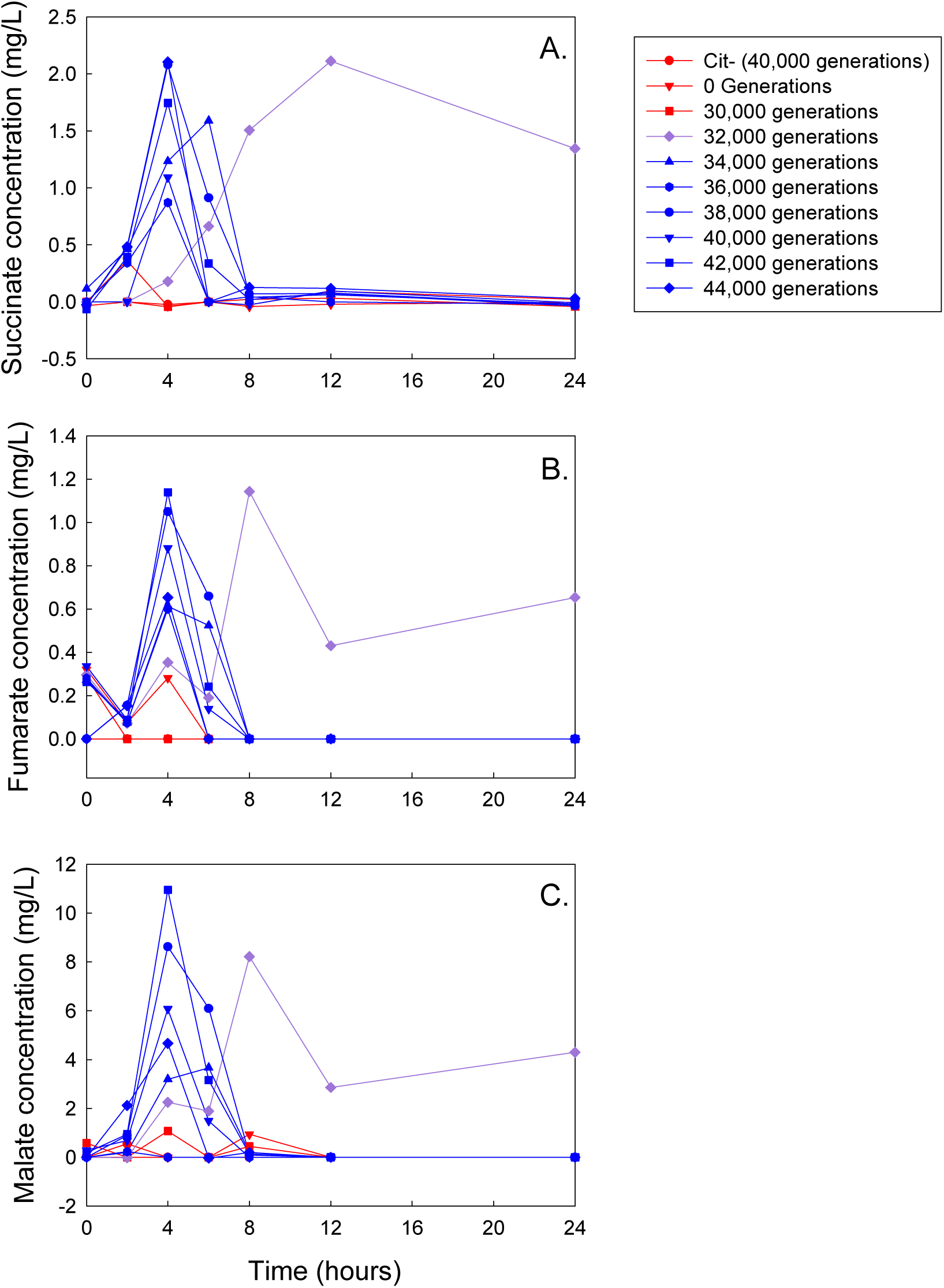
Concentrations of succinate (A), malate (B), and fumarate (C) in filtered medium sampled over a 24-h transfer cycle. Red points and lines show clones with the Cit^−^ phenotype, purple shows a weakly Cit^+^ clone, and blue shows strongly Cit^+^ clones.

The peak succinate levels in medium with Cit^+^ clones from 34,000 generations and later occurred at 4 or 6 h after inoculation and ranged from 0.9 to 2.1 mg/L (Fig. 1A). Succinate concentrations then declined to below detectable levels by 8 h. The culture medium of the 32,000-generation Cit^+^ clone, which grows weakly on citrate, showed a very different pattern, with a peak succinate concentration of 2.1 mg/L at 12 h and 1.3 mg/L remaining at 24 h.

Fumarate and malate exhibited similar patterns to succinate, with peak fumarate concentrations ranging from 0.6 to 1.1 mg/L and peak malate concentrations from 3.2 to 11.0 mg/L (Fig. 1B-C). (The 36,000-generation Cit^+^ clone was atypical, however, in that it did not produce any measurable amount of malate.) As with succinate, the 32,000-generation, weakly Cit^+^ clone released fumarate and malate, but did not fully consume them from the medium. None of the C_4_-dicarboxylates showed any obvious trend, upward or downward, in peak concentration over evolutionary time.

Fig. 2 shows the concentration of citrate in the culture medium during the growth of several Cit^+^ clones. The earliest Cit^+^ clone from generation 32,000 drew down citrate only slightly, if at all, over the full 24-h transfer cycle. By contrast, both Cit^+^ clones that we tested from generations 34,000 and 40,000 rapidly removed citrate in the period from 4 to 6 h, which coincides with the period when the C_4_-dicarboxylates reached their highest levels (Fig. 1). By 9 h, both these clones drew down the citrate concentration to below the detection limit.

**Figure 2.**
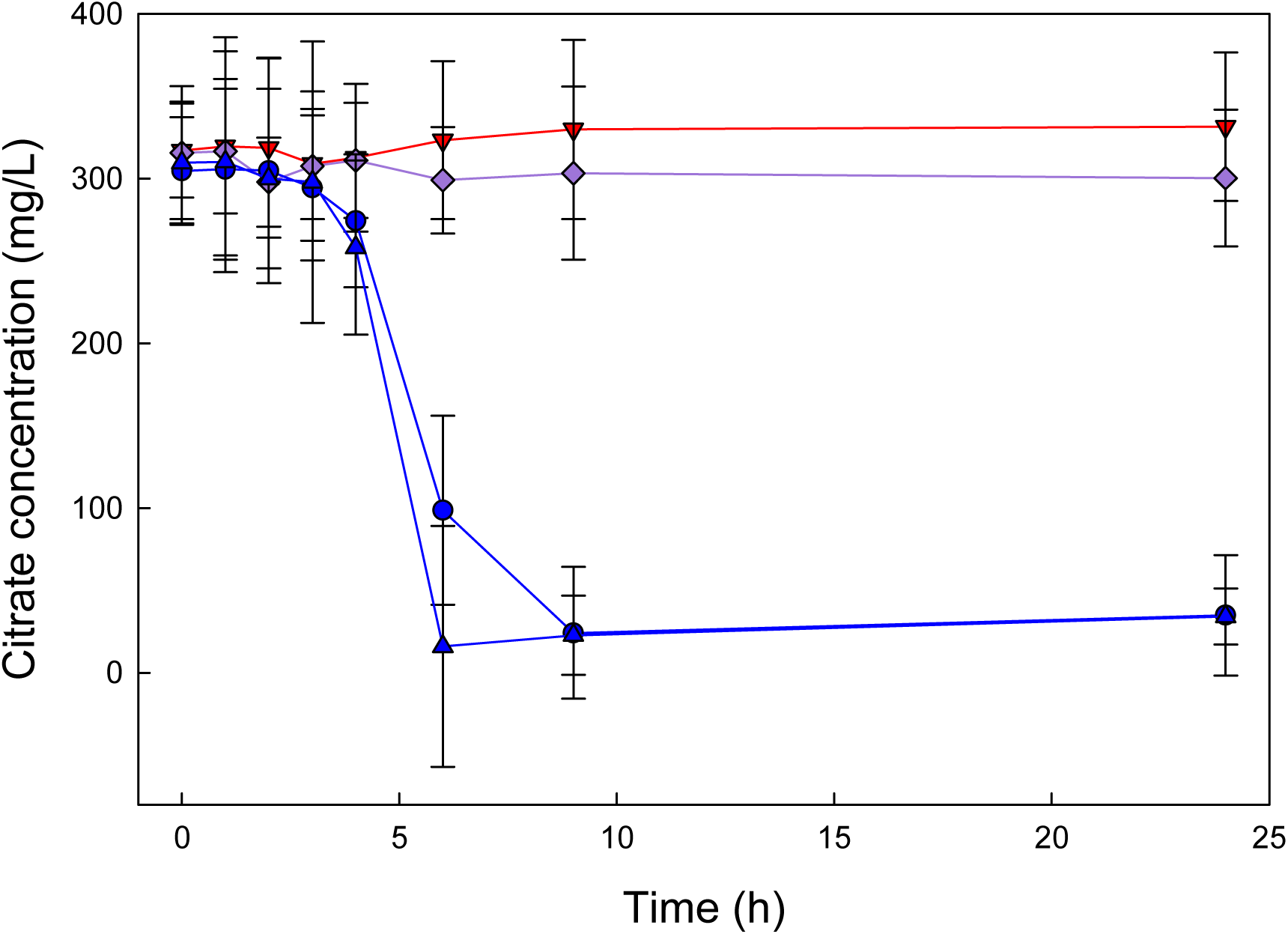
Citrate concentration in filtered medium sampled over a 24-h transfer cycle. Red points and lines show a Cit^−^ clone, purple shows a weakly Cit^+^ clone, and blue shows two strongly Cit^+^ clones. Error bars are 95% confidence intervals.

### Growth on C_4_-dicarboxylates

Clones from the Cit^−^ clade isolated before the demographic shift (30,000–33,000 generations) did not exhibit a measurable increase in OD even after 24 h in glucose-free medium supplemented with any of the three C_4_-dicarboxylates (Fig. 3A-C). However, all Cit^−^ clones from 34,000 generations and later showed improved growth on succinate, fumarate, and malate with OD values significantly greater than zero after 24 h (Fig. 3D-F). These later clones also reached stationary phase in fewer than 24 h, and the time they took to reach stationary phase declined between 34,000 and 43,000 generations (Fig. 3D-F).

**Figure 3.**
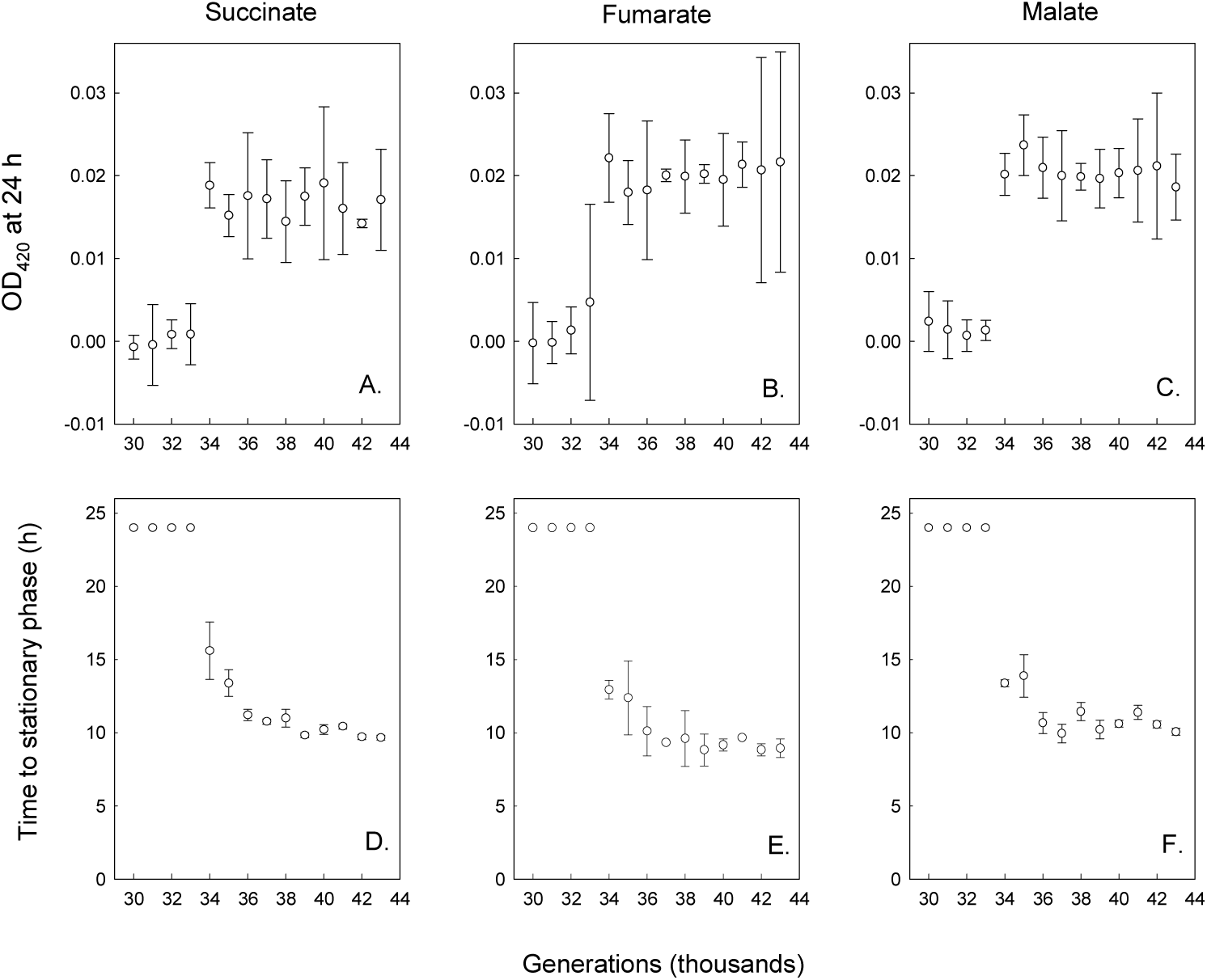
Final OD at 420 nm (A-C) and time to stationary phase (D-F) of clones isolated from the Cit^−^ clade at the various generations indicated grown in DM medium supplemented with 30.5 mg/L succinate, 39.5 mg/L fumarate, or 45.7 mg/L malate. Error bars are 95% confidence intervals.

Clones from the Cit^+^ clade also showed negligible growth on C_4_-dicarboxylates through 32,000 generations (Fig. 4A-C). Beginning at 33,000 generations, Cit^+^ clones had significantly improved growth on succinate, fumarate, and malate as sole carbon sources (Fig. 4A-C). The 32,000-generation clone had a weak Cit^+^ phenotype, but it showed no growth on succinate, fumarate, or malate, indicating that the ability to grow on these carbon sources was not simply a pleiotropic effect of the same mutations that allowed growth on citrate. Unlike the clones from the Cit^−^ clade, Cit^+^ clones did not show consistent improvement across the generations in their growth in succinate medium. For example, the Cit^+^ clones from generations 35,000, 38,000 and 46,000 had lower final OD values than other late-generation Cit^+^ clones and did not reach stationary phase within 24 h when growing on succinate (Fig. 4D). Growth of the Cit^+^ clones on fumarate and malate was similarly variable (Fig. 4E-F), but the particular clones with slower growth or lower final density differed across the three substrates.

**Figure 4.**
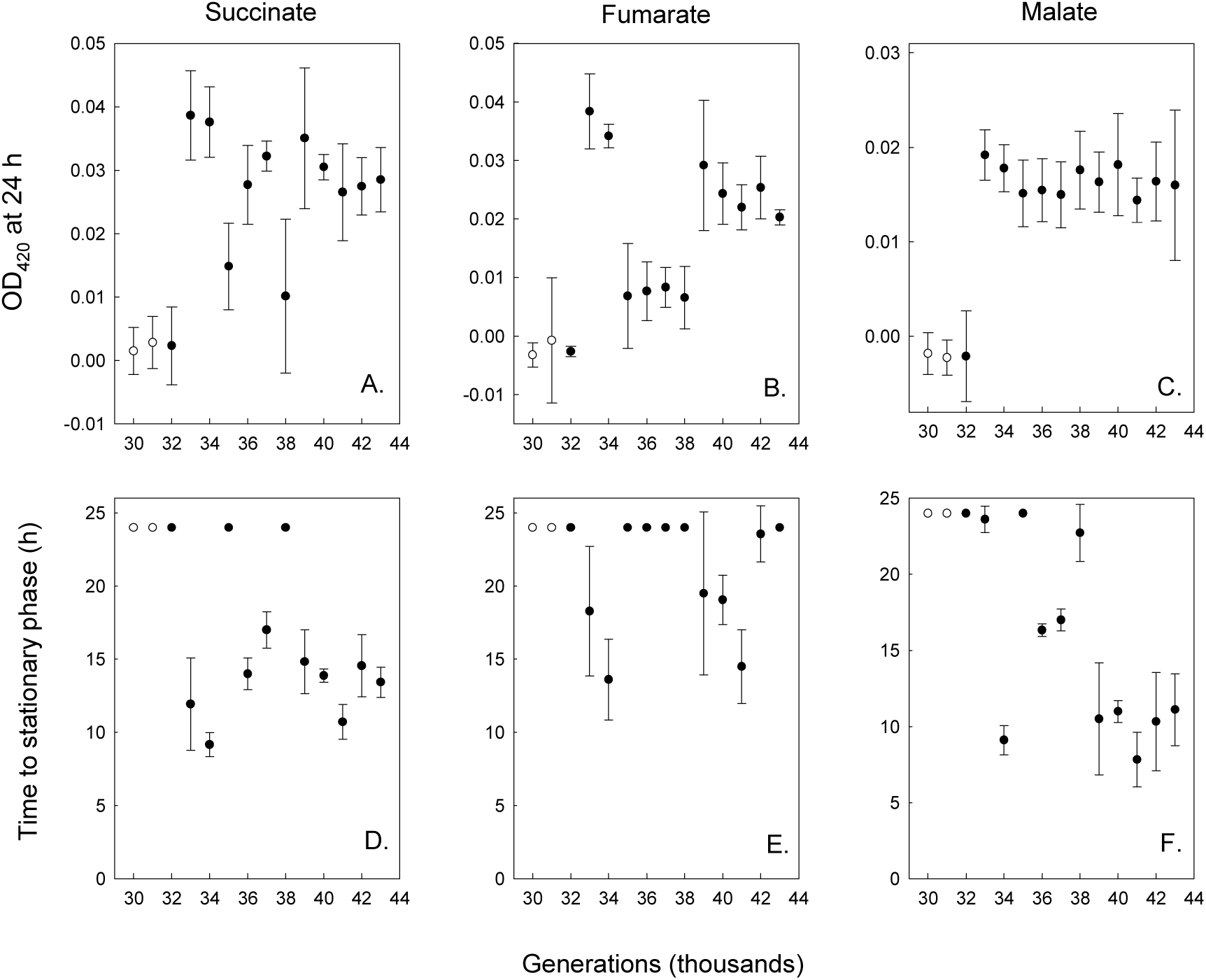
Final OD at 420 nm (A-C) and time to stationary phase (D-F) of clones isolated from the Cit^+^ clade at the various generations indicated grown in M9 medium supplemented with 30.5 mg/L succinate, 39.5 mg/L fumarate, or 45.7 mg/L malate. Open and filled circles show clones with Cit^−^ and Cit^+^ phenotypes. Error bars are 95% confidence intervals.

In contrast to the pattern of improved growth on the C_4_-dicarboxylates succinate, fumarate, and malate after the demographic expansion of the Cit^+^ lineage, clones from the Cit^−^ clade showed reduced growth on acetate, both in terms of final OD values (Fig. 5A) and time to stationary phase (Fig. 5B). Thus, acetate cross-feeding is unlikely to contribute to the coexistence of the Cit^−^ and Cit^+^ lineages. Moreover, the reduced growth on acetate indicates that the improved growth of Cit^−^ clones on C_4_-dicarboxylates is specific to those molecules, rather than a general improvement in either growth or the ability of Cit^−^ cells to shift from growing on glucose to other carbon resources.

**Figure 5.**
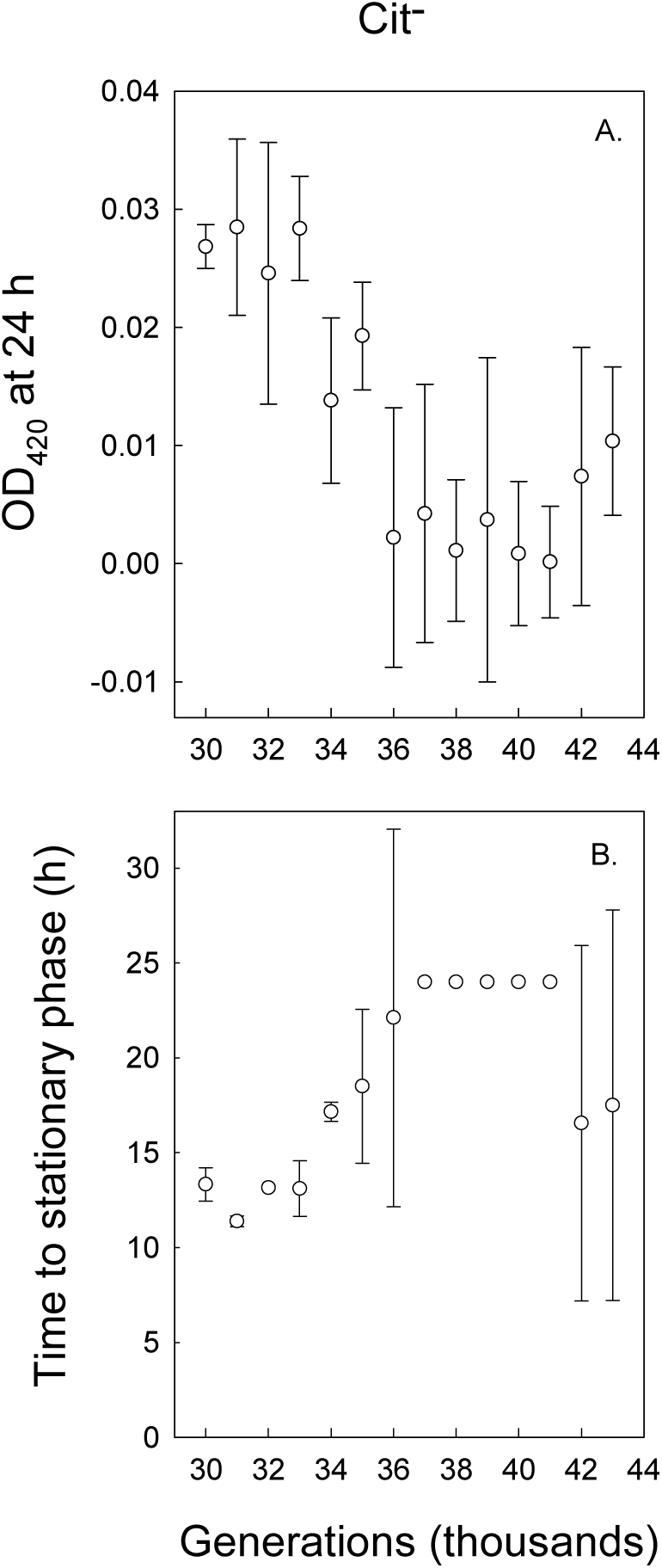
Final OD at 420 nm (A) and time to stationary phase (B) of clones from the Cit^−^ clade growing in DM medium supplemented with 30.5 mg/L acetate. Error bars are 95% confidence intervals.

### Fitness assays

If the Cit^−^ lineage has evolved to feed on byproducts secreted by the Cit^+^ lineage, then it should show evidence of evolving higher fitness, relative to its Cit^−^ predecessors, in the presence of Cit^+^ cells than in their absence. We tested this prediction by competing clones from the Cit^−^ clade isolated at generations 30,000 and 32,000, and then every 1,000 generations from 33,000 to 43,000 generations, against a reference Cit^−^ clone from 30,000 generations, both in the presence and absence of a common Cit^+^ clone that could be excluded when counting the two Cit^−^ competitors (see Methods). In the presence of Cit^+^ cells, the fitness of the Cit^−^ clones trended higher in later generations, with mean fitness values relative to the 30,000-generation reference significantly greater than unity for 9 of the 11 time points tested from generations 33,000 to 43,000 (Fig. 7). However, in the absence of Cit^+^ cells, the fitness of the Cit^−^ clones actually declined over time (Fig. 6). These results not only support the cross-feeding hypothesis, but they also suggest that the adaptation of the Cit^−^ lineage to the cross-feeding niche imposed a tradeoff with respect to growth in its previous glucose-only niche.

**Figure 6.**
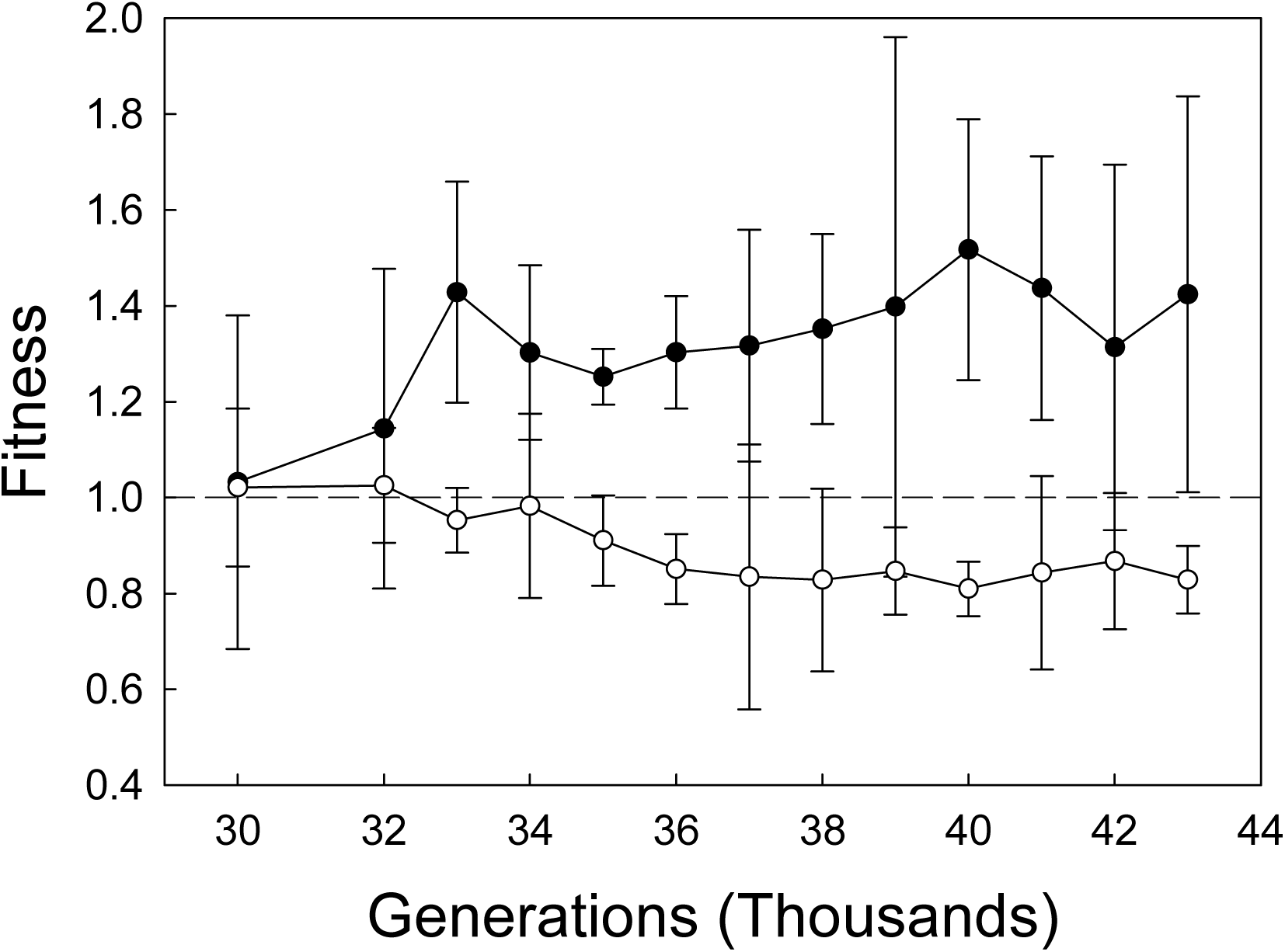
Fitness of clones isolated from the Cit^−^ clade at the various generations indicated relative to a 30,000-generation clone from the Cit^−^ clade. Filled and open symbols indicate competitions between the Cit^−^ clones that were performed in the presence and absence, respectively, of a 40,000-generation Cit^+^ clone. Error bars are 95% confidence intervals.

**Figure 7.**
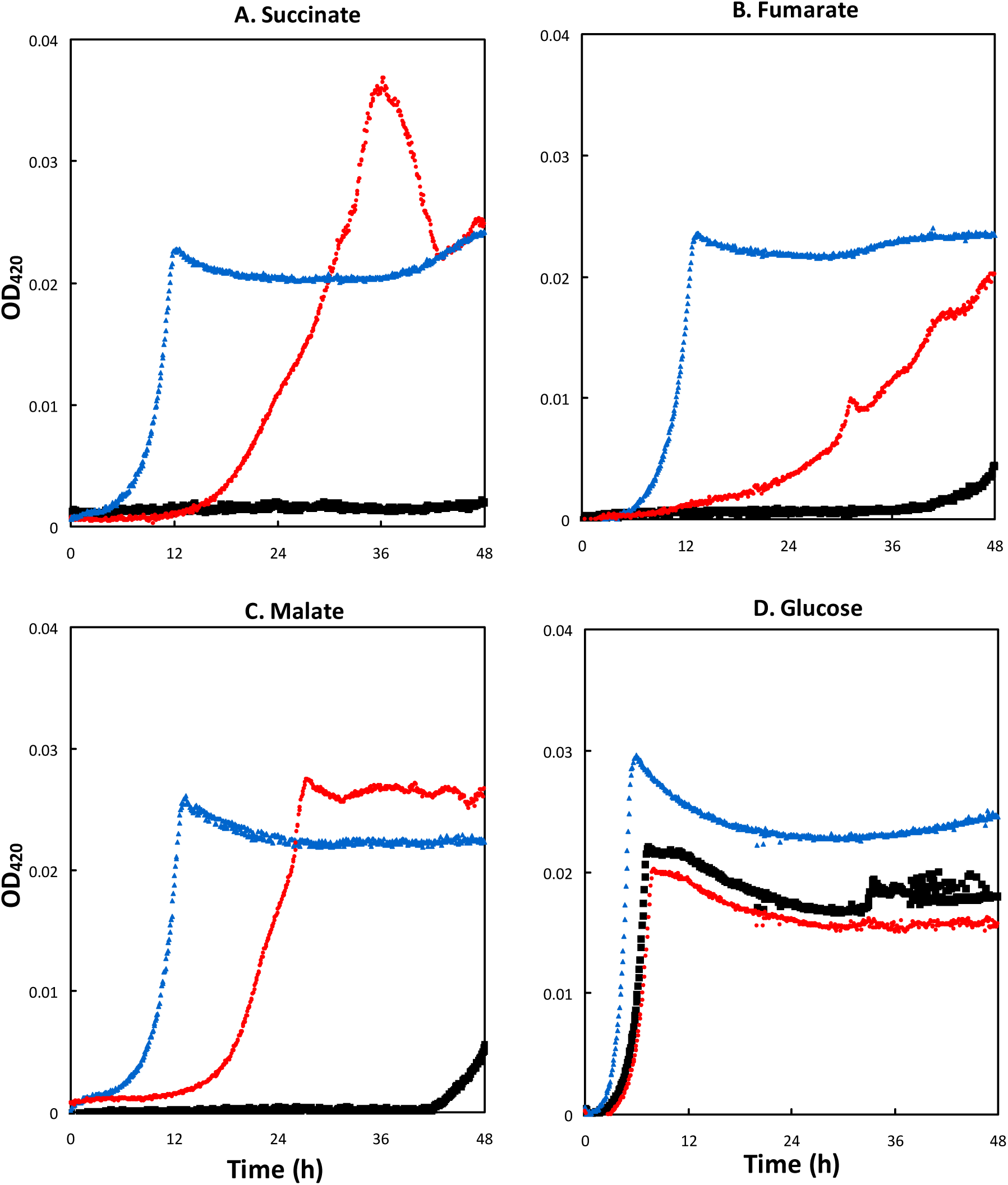
Growth curves showing OD at 420 nm of the ancestral strain (REL606, black squares), a 34,000-generation Cit^−^ clone (ZDB86, blue triangles), and a modified ancestral strain with the evolved *dcuS* allele (ZDB1052, red circles) grown in DM medium supplemented with (A) 30.5 mg/L

Although the overall trend in these data is consistent with our expectations, it is surprising that the increase in the fitness of the Cit^−^ clones, when assayed in the presence of Cit^+^ cells, seems to have begun by 33,000 generations (Fig. 6), whereas we did not see any evidence of their growth on C_4_-dicarboxylates until generation 34,000 and later (Fig. 3). If the differences in fitness of the Cit^−^ cells in the presence and absence of Cit^+^ cells were caused by the evolution of the capacity for growth on the C_4_-dicarboxylates, then we would expect the 34,000-generation C_4_-dicarboxylate-consuming Cit^−^ clone to be more fit than the 33,000-generation Cit^−^ clone in the presence of Cit^+^ cells and, perhaps, less fit in the absence of Cit^+^ cells. This discrepancy might support a more complicated scenario than that envisioned in our hypothesis, or it might indicate limited statistical resolution given only three competition assays performed for each combination of clone and treatment. To gain better resolution of changes in fitness over this critical period, we competed the Cit^−^ clones from 33,000 and 34,000-generations against one another, using a neutral genetic marker to distinguish them (see Methods and Table S1). Consistent with our predictions, the later C_4_-dicarboxylate-consuming clone had a fitness of 1.14 ± 0.04 (mean ± 95% confidence interval, n = 10) relative to the earlier clone that could not use those substrates, when they competed in the presence of Cit^+^ cells. Moreover, in the absence of the Cit^+^ cells, the 34,000-generation Cit^−^ clone had a fitness of only 0.94 ± 0.02 relative to the 33,000-generation Cit^−^ clone, providing further support for the hypothesis that adaptation of the Cit^−^ cells to growth on C_4_-dicarboxylates imposed a tradeoff with respect to their performance in their original glucose-only niche.

### Genomic analysis

We compared the sequenced genomes of seven Cit^−^ clones that differed in their capacity for growth on C_4_-dicarboxylates, generating a list of 19 distinguishing mutations that might underlie that phenotype (Table S2). We performed Sanger sequencing to determine the presence or absence of many of these mutations in six additional Cit^−^ clones: CZB193, CZB194, and CZB195, which did not grow on C_4_-dicarboxylates; and ZDB86, ZDB88, and ZDB92, which did grow on those substrates. These additional data narrowed the list of candidates to nine mutations (Table S2). Of all these candidates, the 5-bp deletion in the *dcuS* gene is the only one that has clear relevance to the C_4_-dicarboxylate growth phenotype. The DcuS protein is a C_4_-dicarboxylate sensor that induces expression of the *dctA* gene, which in turn encodes DctA, a dicarboxylic-acid transporter protein (Janausch et al. 2004). The founding strain of the LTEE had a 5-bp insertion in *dcuS* (Jeong et al. 2009), which eliminated the DcuS function and thereby prevented expression of the *dctA* gene and its encoded dicarboxylic-acid transporter (Yoon et al. 2012; Quandt et al. 2014).

### Genetic manipulation

We tested whether the deletion in *dcuS* was, in fact, a gain-of-function mutation that conferred on the Cit^−^ lineage the ability to grow on the C_4_-dicarboxylates that were secreted by the Cit^+^ lineage. To do so, we constructed ZDB1052, an isogenic derivative of the ancestor REL606 except with the *dcuS* allele from a 34,000-generation Cit^−^ clone called ZDB86. We then measured the growth over 48 h of REL606, ZDB86, and ZDB1052 on succinate, fumarate, and malate.

ZDB1052 had improved growth on succinate, fumarate, and malate compared to REL606 (Fig. 7A-C). The densities achieved by ZDB1052 on these substrates after 48 h were comparable to those reached by ZDB86, the earliest Cit^−^ clone in our study to exhibit the C_4_-dicarboxylate growth phenotype and the source of the evolved *dcuS* allele. However, ZDB1052 grew more slowly on the C_4_-dicarboxylates than did ZDB86. ZDB1052 also grew more slowly on glucose, and reached a slightly lower final density on that substrate, than did REL606, its counterpart with the ancestral *dcuS* allele (Fig. 7D). This result provides direct evidence that adaptation to growth on the C_4_-dicarboxylates caused a tradeoff in terms of growth on glucose, the sole carbon source that was available to the ancestral bacteria. ZDB86 does not show any indication of that tradeoff (Fig. 7A-D), but that is not surprising because it contains many mutations that were beneficial for growth on glucose, which had fixed before citrate utilization evolved and C_4_-dicarboxylates appeared.

The ancestral strain, REL606, did not grow appreciably on C_4_-dicarboxylates during the 24-h interval between transfers of the LTEE, but it did show slight growth between 36 and 48 h on two of these compounds (Fig. 7A-C). This pattern suggests that growth on C_4_-dicarboxylates has a long lag and is slow, but not absent, in the ancestral genotype. Alternatively, this growth could have resulted from a mutant with the ability to grow on C_4_-dicarboxylates. We did not try to distinguish these alternatives because the purpose of collecting these data was to test whether the evolved *dcuS* allele led to improved growth on C_4_-dicarboxylates, which it clearly did.

## Discussion

The ability to grow aerobically on citrate evolved in only one of the twelve LTEE populations (Blount et al. 2008), even after 60,000 generations and billions of spontaneous mutations in each population (Lenski 2004). This evolutionary innovation provided access to a previously unavailable resource, and it caused dramatic ecological changes in the flask-based ecosystem where it arose. Owing to the abundant citrate in the medium, the total population experienced a several-fold increase in density when the Cit^+^ cells became dominant in the population. However, the Cit^+^ lineage did not eliminate all of the non-citrate consumers from the population. Instead, two ecotypes, one Cit^+^ and one Cit^−^, coexisted for thousands of generations in a frequency-dependent manner. We sought to understand the ecological dimensions of this prolonged coexistence. Based on several lines of evidence, we discovered that the Cit^−^ lineage was able to persist, at least in part, by evolving a cross-feeding relationship with the Cit^+^ lineage, one in which three C_4_-dicarboxylates were the secreted metabolites.

We ruled out the possibility that the Cit^−^ lineage coexisted with the Cit^+^ lineage by improvements in their growth on the exogenously supplied glucose. Although Cit^−^ cells initially had an advantage over Cit^+^ cells in competition for glucose (Blount et al. 2008), improved growth on glucose did not sustain the Cit^−^ lineage. In fact, the fitness of the Cit^−^ lineage in the standard glucose-limited medium, when measured in the absence of a Cit^+^ population, actually declined over evolutionary time (Fig. 6).

However, the fitness of clones from the Cit^−^ clade increased over evolutionary time when they competed against their predecessor in the presence of Cit^+^ cells (Fig. 6). This finding supports our hypothesis that the Cit^−^ lineage adapted to changes in the environment caused by the activity of Cit^+^ cells. In particular, Cit^+^ cells secreted the C_4_-dicarboxylates succinate, fumarate, and malate into the culture medium, whereas Cit^−^ cells (including predecessors from the clade that generated the Cit^+^ lineage) did not (Fig. 1). The earliest Cit^+^ clone that we tested showed only weak growth on citrate (Blount et al. 2012), but it, too, released C_4_-dicarboxylates into the medium. However, it did not fully draw them back down during the course of the 24-h transfer cycle, whereas Cit^+^ clones from later generations both released these C_4_-dicarboxylates and then drew them down below detectable levels. Although the observed concentrations of the endogenously produced succinate, fumarate, and malate were low relative to the exogenously added concentrations of glucose and citrate, the total amounts of C_4_-dicarboxylate secreted and available to support growth would probably be much higher if they could be integrated over the course of an entire cycle. In fact, the citrate transporter protein CitT transports one molecule out of the cell for each molecule that it transports into the cell (Pos et al. 1998). While some of these antiporter events may have involved the exchange of one citrate molecule for another, the Cit^+^ populations (except in their earliest form) eventually consumed all of the citrate that was available each day (Fig. 2). It follows logically, then, that the total quantity of C_4_-dicarboxylate molecules released into the medium was equal to the number of citrate molecules consumed, implying substantial opportunity for growth on C_4_-dicarboxylates and, consequently, strong selection to improve growth on those resources. C_4_-dicarboxylate concentrations were generally highest from about 4 to 6 h after inoculation, which coincided with the period when citrate was being drawn down from the medium (Fig. 2), corroborating the biochemical evidence (Pos et al. 1998) that the release of C_4_-dicarboxylates is simultaneous to citrate uptake.

In addition to showing that the Cit^+^ population released C_4_-dicarboxylates into the medium, and that the Cit^−^ population was becoming more fit in the presence (but not the absence) of the Cit^+^ population, we also tested explicitly whether the Cit^−^ lineage evolved to exploit those C_4_-dicarboxylates. Indeed, we found that C_4_-dicarboxylate consumption by the Cit^−^ lineage evolved in close temporal proximity to the rise of the Cit^+^ population to numerical dominance between 33,000 and 33,500 generations. Clones isolated from the Cit^−^ lineage through 33,000 generations showed no measurable growth on succinate, fumarate, or malate, but clones from 34,000 generations and later grew well on all three C_4_-dicarboxylates (Fig. 3). Moreover, the time for populations to reach stationary phase, which is a function of both the length of the lag phase and the subsequent growth rate, declined in later generations, indicating continued adaptation of the Cit^−^ population to growth on C_4_-dicarboxylates. This improvement was not a generalized improvement in fitness because the growth of the Cit^−^ clones on both glucose and acetate declined over this period.

The Cit^+^ lineage also evolved improved growth on C_4_-dicarboxylates, although that improvement was somewhat more sporadic than the improvements seen in the Cit^−^ lineage (Fig. 4). The earliest clone able to grow on citrate, from generation 32,000, did not show any detectable growth on C_4_-dicarboxylates, indicating that improved growth on these compounds was not simply a pleiotropic effect of the ability to consume citrate. The greater variability of growth on C_4_-dicarboxylates in the Cit^+^ lineage suggests that selection to use them was either weaker or less effective in this lineage than in the Cit^−^ lineage. That variability may reflect the fact that Cit^+^ cells can also metabolize citrate, a more abundant and energetically valuable resource than any of the C_4_-dicarboxylates, so that mutations that improve growth on citrate might be favored even if they reduce growth on one or more of these byproducts.

In both the Cit^+^ and Cit^−^ lineages, the mechanism underlying the improved growth on C_4_-dicarboxylates appears to be increased expression of the DctA transporter protein. DctA is a proton motive force-driven transporter that *E. coli* requires for transport of C_4_-dicarboxylates during aerobic metabolism (Davies et al. 1999). The ancestral strain of the LTEE had a 5-bp frameshift mutation in the *dcuS* gene, which encodes a regulator that promotes expression of DctA (Jeong et al. 2009; Quandt et al. 2014). In the Cit^+^ lineage, previous work showed that a mutation in the promoter region of the *dctA* gene caused increased DctA expression and restored the ability to grow on C_4_-dicarboxylates (Quandt et al. 2014). Similarly, in the Cit^−^ lineage, we have showed that a 5-bp deletion in *dcuS* that restored its reading frame enabled growth on C_4_-dicarbolyates (Fig. 7).

The consumption by Cit^−^ cells of the C_4_-dicarboxylates released by Cit^+^ cells is an example of cross-feeding, a phenomenon that has been observed in several previous microbial experiments (Rosenzweig et al. 1994; Treves et al. 1998). Cross-feeding typically occurs when one organism evolves to grow faster by partially degrading a primary resource, leaving a secondary resource for another organism to consume. In this canonical scenario, incomplete degradation of the primary resource is adaptive because it leads to faster growth on the primary resource (Doebeli 2002; Pfeiffer and Bonhoeffer 2004). However, the evolution of cross-feeding between the Cit^+^ and Cit^−^ lineages follows a somewhat different pattern. The release of C_4_-dicarboxylates by the Cit^+^ cells does not itself appear to be beneficial to them. Indeed, while the Cit^−^ lineage evolved increased growth on the C_4_-dicarboxylates, so too did the Cit^+^ lineage. Thus, unlike the typical cross-feeding relationship, the two ecotypes have evolved increased, not decreased, overlap in their resource usage. Instead of being an adaptive trait, cross-feeding in this case appears to reflect evolutionary ‘bricolage’ (Jacob 1977), whereby evolution tinkers with existing components to solve new problems.

The Cit^+^ lineage studied here evolved the ability to consume citrate via multiple mutations including, in particular, a complex rearrangement that allowed aerobic expression of an existing CitT transporter (Blount et al. 2012). Most *E. coli* cells can express the CitT protein only under anaerobic conditions, where cells ferment citrate into succinate (but only when they also have another growth substrate in addition to citrate). Under anaerobic conditions, the use of a transporter that exchanges citrate for succinate has a clear benefit because succinate is an end product of fermentation. However, in the Cit^+^ lineage, the CitT transporter is expressed under carbon-limited, aerobic conditions where succinate and other C_4_-dicarboxylates provide a useful source of energy and carbon. Many other bacterial species use proton or sodium gradients for citrate transport (Boorsma et al. 1996; Bott 1997). Furthermore, in the only other reported case where mutations gave rise to aerobic consumption of citrate by *E. coli* (Hall 1982), the citrate importation appears to have been powered by the proton motive force (Reynolds and Silver 1983), although the genetic and mechanistic bases are unknown in that case. If the Cit^+^ ecotype in the LTEE had instead evolved to use a proton or sodium gradient for citrate uptake, then it would not have exported C_4_-dicarboxylates. A hypothetical Cit^+^ mutant that employed a proton or sodium gradient would probably be favored over the evolved Cit^+^ bacteria that release valuable molecules that are often lost to competitors. Instead, as a consequence of evolutionary tinkering, or bricolage, the Cit^+^ lineage first evolved to express the previously unexpressed CitT antiporter under aerobic conditions (Blount et al. 2012) and then to reabsorb the exported C_4_-dicarboxylates via the proton motive force by using the previously unexpressed DctA transporter (Quandt et al. 2014). While this solution seems not to have been the most elegant or efficient, it nonetheless proved highly successful for the Cit^+^ clade. At the same time, this solution provided an opportunity for the Cit^−^ lineage to expand its niche as well, transitioning from being a glucose specialist to also consuming some of the C_4_-dicarboxylates released by the Cit^+^ cells.

## Acknowledgments

We thank Justin Meyer for assistance with λ phage; Neerja Hajela for laboratory assistance; Mike Wiser, Alita Burmeister, and Rohan Maddamsetti for helpful comments on this paper; and the technical staff at the MSU Mass Spectrometry Core Facility for assistance with GCMS. This research was supported, in part, by EPA STAR and MSU Distinguished Graduate Student Fellowships to C.B.T., an NSF grant (DEB-1019989) to R.E.L., a grant from the John Templeton Foundation to R.E.L. and Z.D.B., and the BEACON Center for the Study of Evolution in Action (NSF Cooperative Agreement DBI-0934).

